# Computer simulations reveal pathogenicity and inheritance modes of hearing loss-causing germinal variants

**DOI:** 10.1101/2022.05.02.490275

**Authors:** Cheng-Yu Tsai, Ying-Chang Lu, Yen-Hui Chan, Yuan-Yu Chang, Shu-Wha Lin, Tien-Chen Liu, Chuan-Jen Hsu, Pei-Lung Chen, Lee-Wei Yang, Chen-Chi Wu

## Abstract

Variants in the gap junction beta-2 (*GJB2*) gene are the most common cause of hereditary hearing impairment. However, how *GJB2* variants lead to local physicochemical and structural changes in the hexameric ion channels of connexin 26 (Cx26), resulting in hearing impairment, remains elusive. In the present study, using molecular dynamics (MD) simulations, we showed that detached inner-wall N-terminal “plugs” aggregated to reduce the channel ion flow in a highly prevalent V37I variant in humans. To examine the predictability of the computational platform, an artificial mutant, V37M, of which the effect was previously unknown in hearing loss, was created. Microsecond simulations showed that homomeric V37M Cx26 hemichannels had an abnormal affinity between the inner edge and N-termini to block the narrower side of the cone-shaped Cx26, while the most stable heteromeric channels did not. Consistent with these predictions, homozygous V37M transgenic mice exhibited apparent hearing loss, but not their heterozygous counterparts, indicating a recessive inheritance mode. Reduced channel conductivity was found in *Gjb2*^V37M/V37M^ outer sulcus cells and Claudius cells but not in *Gjb2*^WT/WT^ cells. We view that the current computational platform could serve as an assessment tool for the pathogenesis and inheritance of *GJB2*-related hearing impairments and other diseases caused by connexin dysfunction.

## 1. Introduction

Gap junctions (GJs), formed by two hemichannels assembled from hexameric connexin subunits, are crucial for electrolyte and metabolite homeostasis in the mammalian inner ear.^[1]^ The connexin 26 (Cx26) channel protein, encoded by gap junction beta-2 (*GJB2*), is the most important connexin in the inner ear. *GJB2* variants are the most common genetic cause of sensorineural hearing impairment (SNHI) in humans, and can be inherited in both autosomal recessive and dominant modes.^[2]^ Expressed in the supporting cells of the organ of Corti, stria vascularis, spiral ligament, and spiral limbus^[3]^ in the cochlea, Cx26 has been implicated in potassium recycling,^[4]^ intercellular signaling,^[5]^ and development of the organ of Corti.^[6]^

To date, over 400 pathogenic *GJB2* variants have been recorded in the Deafness Variation Database (http://deafnessvariationdatabase.org/).^[7]^ Of these, the V37I (c.109G>A) variant is of particular interest because of its high prevalence and enigmatic pathogenicity. This variant is prevalent in the normal Asian population, with an allele frequency of 8.35% (gnomAD, v2.1.1),^[8]^ ranging from ∼1.0% in Japanese^[9]^ and Korean,^[10]^ 4.3% in Thai,^[11]^ and 6.2%–8.9% in Han Chinese populations.^[12]^ At least five million Asian people are estimated to be homozygous for V37I,^[13]^ indicating that V37I might be the single most common deafness-associated variant worldwide. Owing to its high allele frequency, V37I was previously regarded as a benign polymorphism.^[11, 12b]^ However, subsequent studies have indicated that V37I might be a recessive pathogenic variant contributing to mild-to-moderate SNHI with incomplete penetrance.^[2b, 10, 12a, 13-14]^ In a recent longitudinal study, we described the progressive nature of SNHI in subjects homozygous for V37I; however, hearing levels varied remarkably among individuals.^[15]^

Changes in the amino acid residues of Cx26 may confer structural^[16]^ and biochemical effects^[16a, 16b, 17]^ on the gating of gap junction channels. *In silico* molecular dynamics (MD) simulations have been applied to investigate the structural effects of several missense *GJB2* variants, including M34T,^[16c]^ T8M,^[18]^ G12R,^[19]^ and N14K,^[20]^ on the gating mechanisms of the Cx26 hemichannel. Biochemical assays of V37I revealed reduced permeability of gap junctions,^[21]^ suggesting that V37I may exert structural effects implicated in the plug-mediated gating mechanism similar to M34T;^[17c, 22]^ however, the detailed mechanisms underlying V37I-mediated reduced permeability remain unclear. Spatially, residue 37 (Res 37) is located in the middle of transmembrane domain 1 (TM1) and adjacent to the N-terminal helices (NTHs), wherein the TM1-loop-NTH domain constitutes the major component of the inner edge of the pore. Therefore, it is of interest to clarify how V37I exerts its effects on Res 37 and leads to conformational changes in Cx26.

MD simulations can potentially predict the functionality of other missense variants of Cx26. To test this hypothesis, an *in silico* model was constructed using a mouse-based Cx26 homolog with a novel mutant at Res 37, namely V37M, in homomeric and heteromeric arrangements. The clinical data for the mutant were not available to interfere with or precondition our MD-based prediction. Our simulations showed normal functional Cx26 channels for *Gjb2*^WT/WT^ and *Gjb2*^WT/V37M^, but not *Gjb2*^V37M/V37M^, which was confirmed by hearing loss/noise vulnerability tests in homozygous V37M transgenic mice. The reduced molecular conductivity of *Gjb2*^V37M/V37M^ was confirmed using *in vitro* dye transport assays.

## 2. Results

### 2.1. V37I in human *in silico* Cx26 hemichannel revealed reduced pore volume and potassium permeability

To investigate the abnormal gating mechanisms influenced by the missense variant V37I, a standard *in silico* model based on the crystal structure of human Cx26 hemichannel (hereafter called “hCx26”; PDBID: 2ZW3^[23]^) was first constructed, which was embedded into the 1,2-dioleoyl-sn-glycero-3-phosphocholine (DOPC) bilayer and solvated under the physiological condition that mimicked the endolymph of the cochlea (**Figure 1**a; “WT-hCx26”), as well as its mutant with homomeric V37I variant (“V37I-hCx26”) (Figure 1b).

**Figure 1.**
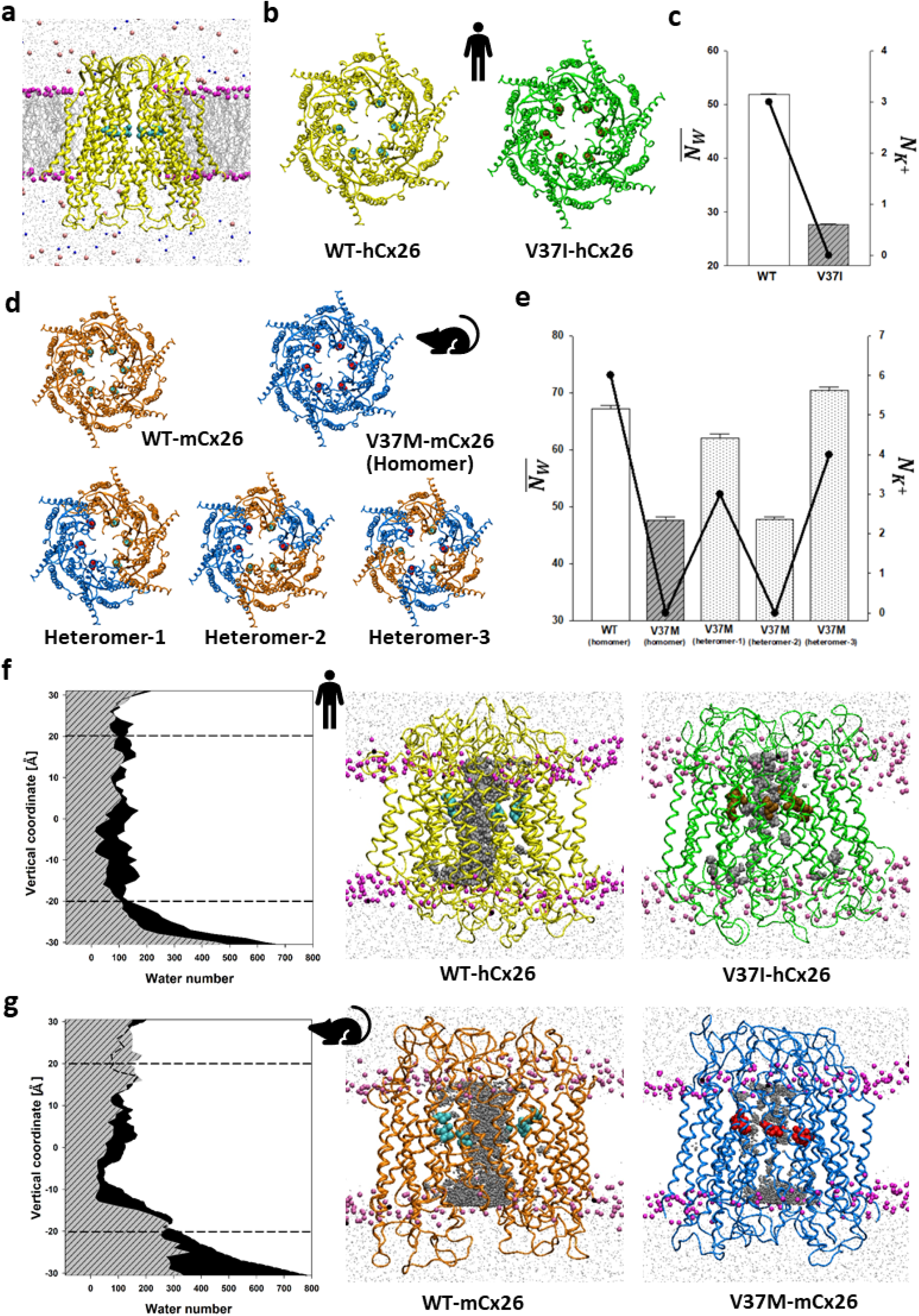
Overview of the membrane-embedded Cx26 hemichannel and changes in pore volume, potassium ion permeability, and water channel caused by Res37 variants in human and mouse Cx26 models. (a) Side view of the whole Cx26-DOPC model containing the human wild-type Cx26 hemichannel [WT-hCx26 (yellow) with hexameric Val37 (cyan)] buried in DOPC bilayers (violet beads are choline groups) and solvated with 158 mM potassium (indigo) and chloride (pink) ions. (b and d) Top views of WT-hCx26 and its mutant with homomeric Ile37 [V37I-hCx26 (green) with hexameric Ile37 (ochre)] as well as the homologous mouse model, including the wild type [WT-mCx26 (orange) with hexameric Val37 (cyan)] and its mutants with homomeric Met37 [V37M-mCx26 (blue) with hexameric Met37 (red)] and heteromeric counterparts (heteromers-1 to −3 of V37M-mCx26 in equal ratio of 3:3 under different arrangements of WT and V37M monomers). (c and e) Mixed graphs containing average water numbers inside the hemichannel pore 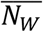 illustrated by bar plots on the left axis) and total potassium ions passing through the hemichannels 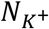 illustrated by line charts on the right axis) for hCx26 and mCx26 models, respectively. (f and g) Left panel: area plots of accumulated water distribution in wild-type (in black) and homomeric mutants (slash-filled gray) of hCx26 and mCx26 models, respectively, during the equilibrated 4 μs simulations along the membrane normal (transmembrane helix) of the waterbox. Dashed lines represent the average coordinates of choline groups along the vertical axis in the upper and lower leaflets of the DOPC membrane; right panel: water channel including the accumulated trajectories of water (in gray) during the equilibrated 4 μs of molecular dynamic (MD) simulations as well as the corresponding Cx26-DOPC conformer at the final snapshot of MD simulation for the wild type and mutants with homomeric Res37 variants of hCx26 and mCx26 models, respectively.

Four microsecond-long MD simulations were performed to investigate the conformations of WT-hCx26 and V37I-hCx26 using three metrics: pore volume (inner channel space), potassium permeability, and retention time of potassium ions. The water channeling of V37I-hCx26, represented by the average water number inside the hemichannel 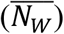, was found reduced as compared with the wild-type (WT vs. mutant was 51.76 ± 0.34 vs. 26.52 ± 0.22 water molecules, per Student’s t-test, *p* < 0.001, see Figure 1c and **Table 1**). As for the potassium permeability, as defined by the number of potassium ions that passed through the pore during the simulations 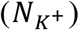, three ions passed through the pore of WT-hCx26, whereas no potassium ions passed through V37I-hCx26 (see line chart of Figure 1c and Table 1). The reduced permeability of V37I-hCx26 was also in line with the prolonged retention time of potassium ions 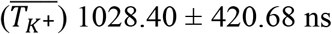 compared to the wild-type 219.31 ± 60.88 ns (Table 1). The difference in potassium permeabilities between WT-hCx26 and V37I-hCx26 is further illustrated in Supplementary videos S1 and S2, respectively. These results indicate that the V37I variant could lead to the dysfunction of human Cx26 through a reduced pore volume, impaired permeability, and prolonged retention time of potassium ions in the hemichannel. The findings from the MD simulations are consistent with the reduced permeability observed in previous biochemical cell line studies of Cx26 with V37I.^[21]^

**Table 1.** Average water number inside the channel 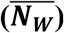, potassium permeability 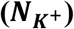 and average retention time of potassium 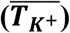 in various hCx26 or mCx26 hemichannels.

| Cx26 hemichannel | $\overline{N_W}$ <sup>a)</sup> | $N_{K^+}$ <sup>b)</sup> | $\overline{T_{K^+}}$ <sup>c)</sup> |
| --- | --- | --- | --- |
| WT-hCx26 | 51.83 ± 0.19 | 3 | 219.31 ± 60.88 |
| V37I-hCx26 | 27.62 ± 0.12 *** | 0 | 1028.40 ± 420.68 ** |
| WT-mCx26 | 67.22 ± 0.60 | 6 | 123.07 ± 25.93 |
| V37M-mCx26 (homomer) | 47.67 ± 0.54 *** | 0 | 289.07 ± 125.7 * |
| V37M-mCx26 heteromer-1 | 62.09 ± 0.68 *** | 3 | 147.44 ± 44.46 |
| V37M-mCx26 heteromer-2 | 47.84 ± 0.35 *** | 0 | 253.22 ± 69.81 * |
| V37M-mCx26 heteromer-3 | 70.50 ± 0.60 *** | 4 | 95.73 ± 17.64 |
| $\sigma_p$ <sup>d)</sup> | | 0.93 | -0.99 |

<sup>a)</sup> Average water number inside the hemichannel during the 4 μs simulation (mean ± SEM; \*\*\*: $p < 0.001$ , as compared to the wild type); <sup>b)</sup> potassium permeability, defined as the frequency of potassium ions passing through the hemichannel during the 4 μs simulation. <sup>c)</sup> Average retention time (in ns) of potassium ions at least 20 ns inside the pore during the simulation (mean ± SEM; \*: $p \leq 0.05$ , \*\*: $p < 0.01$ , as compared to the wild type); <sup>d)</sup> Pearson correlation coefficients of $\overline{N_W}$ and $N_{K^+}$ as well as $\overline{N_W}$ and $\overline{T_{K^+}}$ for various mCx26.

### 2.2. V37M in mouse *in silico* Cx26 hemichannel revealed impaired channeling from homomeric assembly but not from heteromeric counterparts

To further examine the sensitivity of our MD models, another missense variant, V37M, of our research interest, was examined. The variant was not reported in SNHI-related clinical studies previously so as to remain unknown for its influence on the functionality of hemichannels; based on methionine’s side-chain similarity with those of Val/Ile and a slightly higher polarity than both (due to its sulfur atom), our computer model from the present study could be used to identify subtle differences prior to experimental confirmation through animal studies. Irrespective of the lack of an experimentally determined mouse Cx26 structure, a homology structural model of mouse Cx26 (mCx26) as well as its V37M variant (“V37M-mCx26”) were constructed *in silico* by SWISS-MODEL^[24]^ based on the crystal structure of hCx26 (“WT-mCx26”) given the high (93%) amino acid sequence identity shared between hCx26 and mCx26 (Figure 1d). The resulting structures were embedded in the DOPC bilayer. After 4 μs of standard MD simulations at body temperature (310K) and room pressure (1 bar), V37M-mCx26 demonstrated blockage and therefore reduced permeability compared with WT-mCx26, where WT-mCx26 vs. V37M-mCx26 for 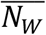 is 67.22 ± 0.6 vs. 47.67 ± 0.54 water molecules, for 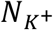 is 6 vs. 0 potassium ions and for 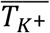 is 123.07 ± 25.93 vs. 289.07 ± 125.7 ns (see Table 1 for definition). As illustrated in Supplementary videos S3 and S4, V37M-mCx26 demonstrated obstruction of potassium ions transport up to 3 μs, which was in line with the reduced pore volume and permeability.

In addition to the homomeric hemichannel, we wondered whether the V37M variant was haploinsufficient to compromise the hemichannel function; to examine this, three *in silico* models with heteromeric V37M mCx26 (called heteromer-1 to −3 of V37M-mCx26) were constructed, which were composed of the same 1:1 ratio of WT and V37M monomers but with different hexameric arrangements (Figure 1d). In 4 μs MD simulations, both heteromer-1 and −3 retained the normal pore volume and retention time of potassium, similar to WT-mCx26 (see Figure 1c and Table 1), while heteromer-2 showed a reduced permeability. However, heteromer-1 and −3 have a lower configurational potential (*E*_conf_) −6.52×10^4^ and −6.50×10^4^ kJ mol^-1^, respectively, than that of heteromer-2 (−6.41×10^4^ kJ mol^-1^), indicating that heteromer-1 and −3 are predominant in conformational population according to Boltzmann relation (probability of a conformation ∼ exp(-*E*_conf_/*k*_*B*_*T*), where *k*_*B*_ and *T* are Boltzmann constant and temperature, respectively).

Overall, our *in silico* mCx26 models demonstrated that V37M could result in dysfunction of the hemichannel in homomers and sufficient function in the predominant heteromers. The pore volume in our mCx26 models also exhibited high correlations with both the potassium permeability (σ_*p*_= 0.93) and retention time of potassium (σ_*p*_ = −0.99), suggesting that the compromised permeability of the hemichannel was strongly associated with the shrunken pore volume.

### 2.3. Reduced water channeling in both V37I-hCx26 and V37M-mCx26 as a function of the depth of hemichannels

To monitor the mutant-caused pore shrinkage longitudinally, the distribution of water molecules inside the pore was monitored by a cumulative count of water molecules in all the snapshots during the last microsecond of the simulation. V37I-hCx26 showed a notable shrinkage in the lower half, 0 to 5Å above the membrane center (where 0Å is at), (Figure 1f), which is below the location of Res37, at average 6.4 Å above the membrane center.

Regarding the mCx26 models, WT-mCx26 demonstrated an hourglass-like water channel, with the narrowest point located toward the cytoplasm end. V37M-mCx26 exhibited an apparent narrowing of the channel from 8 (near the M37 residues) to −4 Å along the membrane normal (Figure 1g). These observations suggest that V37I-hCx26 and V37M-mCx26 may contribute to hemichannel obstruction through different mechanisms.

### 2.4. *Gjb2*^V37M/V37M^ mice demonstrated a faster deterioration of hearing with age

To validate our structural dynamics models, knock-in mouse models were constructed in a C57BL/6 background harboring homozygous and heterozygous *Gjb2* V37M variants (that is, *Gjb2*^*V37M/V37M*^ and *Gjb2*^*WT/V37M*^ mice, respectively). The hearing thresholds were recorded for 13 *Gjb2*^*WT/WT*^, 19 *Gjb2*^*V37M/WT*^, and 30 *Gjb2*^*V37M/V37M*^ mice at 4, 12, 28, and 44 weeks. Consistent with previous studies, C57BL/6 mice developed progressive hearing loss with increasing age.^[25]^ There was no significant difference in hearing thresholds between *Gjb2*^*V37M/WT*^ and *Gjb2*^*WT/WT*^ mice at any frequency and age, whereas *Gjb2*^*V37M/V37M*^ mice demonstrated a faster and significant deterioration of hearing with age compared to *Gjb2*^*WT/WT*^ and *Gjb2*^*V37M/WT*^ mice mainly in 28 and 44 weeks (**Figure 2**a and **Table S1**), indicating that V37M was inherited in an autosomal recessive manner in mice. All three genotypes demonstrated normal appearance of hair cells in the organ of Corti (Figure 2b to 2d) without significant differences in the cochleogram (hair-cell-count graph) at 4 weeks (Figure 2e). These results indicate that the progressive hearing loss in *Gjb2*^*V37M/V37M*^ mice was found without loss of hair cells.

**Figure 2.**
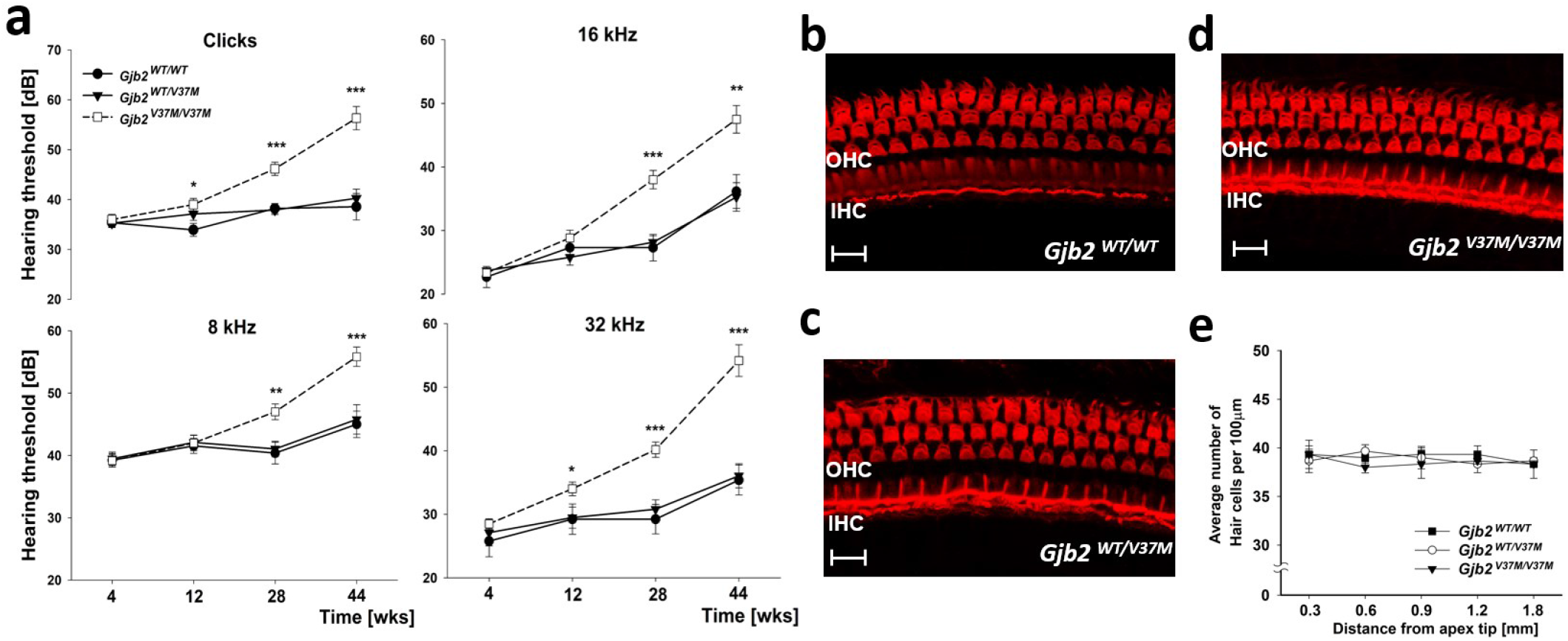
Faster deterioration of hearing loss without significant changes of hair cells in *Gjb2*^*V37M/V37M*^ mice. (a) *Gjb2*^*V37M/V37M*^ mice demonstrated progressive hearing loss with age at a faster deterioration rate in all frequencies than that demonstrated by *Gjb2*^*WT/WT*^ and *Gjb2*^*V37M/WT*^ mice (unit: dB of sound pressure level; mean ± SEM). (b–d) At four weeks, the cochleograms of all three genotypes showed no significant changes of viable outer hair cell numbers in the middle turn of the organ of Corti. (e) The average number of hair cells for each genotype was counted using per high-power field of cochleograms from regions progressively distant from the apex of the cochlea (mean ± SEM). (***, *p* < 0.001; **, *p* < 0.01; *, *p* < 0.05; IHC, inner hair cells; OHC, outer hair cells; Bar = 10 μm)

### 2.5. Normal expression and localization of Cx26 was shown, but compromised GJ-mediated metabolite transfer occurred in V37M mice

To investigate whether the V37M variant affects the expression and trafficking of the protein, the formation of GJs in the supporting cells of the cochlea was examined using immunolabeling for mCx26. GJ plaques, as revealed by the clustering of mCx26 immunoreactivities, were identified in the outer sulcus cells and Claudius cells (**Figure 3**). The sizes of the GJ plaques were similar in *Gjb2*^*WT/WT*^ and *Gjb2*^*V37M/V37M*^ mice (Figure 3a). This observation was confirmed by quantification of the density of mCx26 immunoreactivities using the MetaMorph software: the fluorescence intensity of mCx26 immunoreactivities was 78.2 ± 2.15 pixels in *Gjb2*^*WT/WT*^ mice (n = 3) and 77.2 ± 1.93 pixels in *Gjb2*^*V37M/V37M*^ mice (*n* = 3), showing no difference between the two groups (Figure 3b; Student’s t-test, *p* > 0.05).

**Figure 3.**
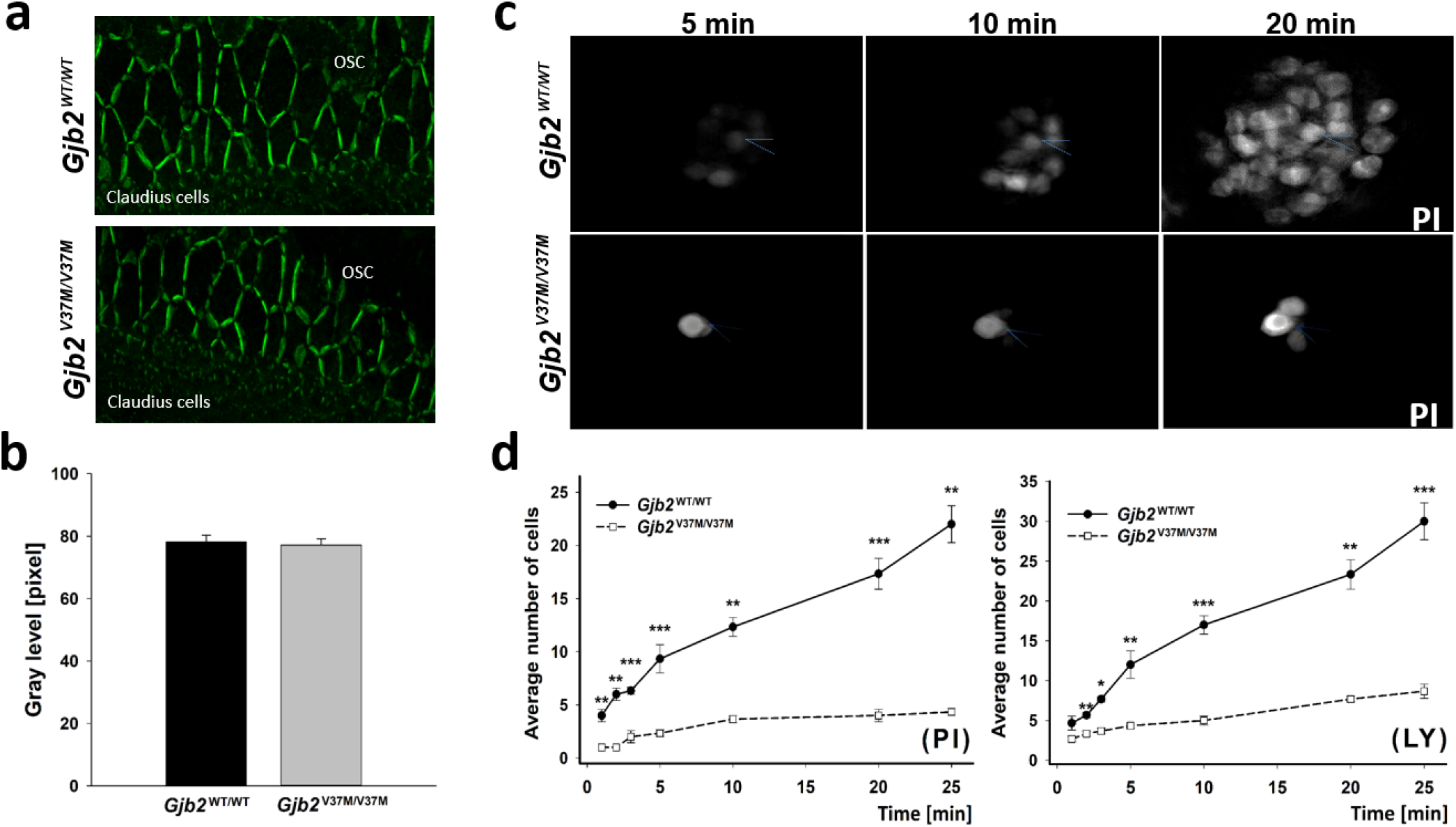
Normal expression of Cx26 but abnormal channel permeability in *Gjb2*^*V37M/V37M*^ mice. (a and b) Patterns of Cx26 immunoreactivities in outer sulcus and Claudius cells of *Gjb2*^*WT/WT*^ and *Gjb2*^*V37M/V37M*^ mice (a) showed no significant difference in the density of Cx26 immunoreactivities between the two strains (b). (OSC, outer sulcus cells; Bar = 5 μm; number of images = 15 [n = 3, five images per mouse for each strain]; density of immunoreactivities are calculated as mean ± SEM). (c) Time-lapse recordings of intercellular dye transfer in *Gjb2*^*WT/WT*^ and *Gjb2*^*V37M/V37M*^ mice after propidium iodide (PI) was injected into a single Claudius cell. Intercellular PI diffusion patterns in *Gjb2*^*WT/WT*^ (upper row panels) and *Gjb2*^*V37M/V37M*^ (lower row panels) mice are demonstrated. Time (in min) after injection of PI is shown in the top left corner of the images. Dotted lines indicate the injection site of needles. (**d**) Comparison of the number of cells (y-axis) receiving fluorescent dyes through GJ-mediated diffusion in *Gjb2*^*WT/WT*^ and *Gjb2*^*V37M/V37M*^ mice. The x-axis indicates the time after the establishment of whole-cell recording conformer. Cells were counted per high-power field, and data are shown as mean ± SEM (n = 5). The number of cells receiving dye transfer in the sensory epithelium of *Gjb2*^*V37M/V37M*^ mice was consistently less at all time points than in *Gjb2*^*WT/WT*^ mice for both the positively charged PI and the negatively charged lucifer yellow (LY). (*: *p* < 0.05; **: *p* < 0.01; ***: *p* < 0.001).

We then investigated whether V37M affected GJ-mediated metabolic coupling in supporting cells using dye diffusion assays in flattened cochlear preparations. Two well-characterized fluorescent dyes, positively charged propidium iodide (PI) and negatively charged lucifer yellow (LY), are known to pass through the cochlear GJs. PI and LY were injected into single Claudius cells^[26]^ to quantify the differences in metabolic coupling among cells. Time-lapse recordings were performed to assess the time course of PI and LY diffusion (Figure 3c). Data obtained from *Gjb2*^*WT/WT*^ and *Gjb2*^*V37M/V37M*^ mice at various time points after the start of dye injection showed that the extent of fluorescent dye diffusion gradually increased with time (Figure 3d). These data were quantified by plotting the number of dye-recipient cells as a function of time after dye injection. This demonstrated that the number of cells undergoing dye transfer in the sensory epithelium of *Gjb2*^*V37M/V37M*^ mice was consistently lower at all time points, indicating that V37M caused a deficit in GJ-mediated metabolite transfer among cells in the sensory epithelium of the cochlea.

### 2.6 Gjb2^V37M/V37M^ mice revealed higher shifts in hearing thresholds after noise exposure

Consistent with our previous report,^[27]^ mice of all genotypes after 3 h noise exposure in the level of 115 dB (unit of Sound Pressure Level, SPL), developed significant elevation of hearing thresholds immediately (30 min) and recovered gradually over 14 days (**Figure 4**a). Notably, *Gjb2*^*V37M/V37M*^ mice (n = 5 for each background) revealed significantly higher shifts in hearing thresholds within the first few days after noise exposure than *Gjb2*^*WT/WT*^ mice. The significant difference in the hearing threshold shift between *Gjb2*^*V37M/V37M*^ and *Gjb2*^*WT/WT*^ mice persisted for longer than 4 days (Figure 4a). Overall, these data indicate that *Gjb2*^*V37M/V37M*^ mice were more vulnerable to noise than *Gjb2*^*WT/WT*^ mice.

**Figure 4.**
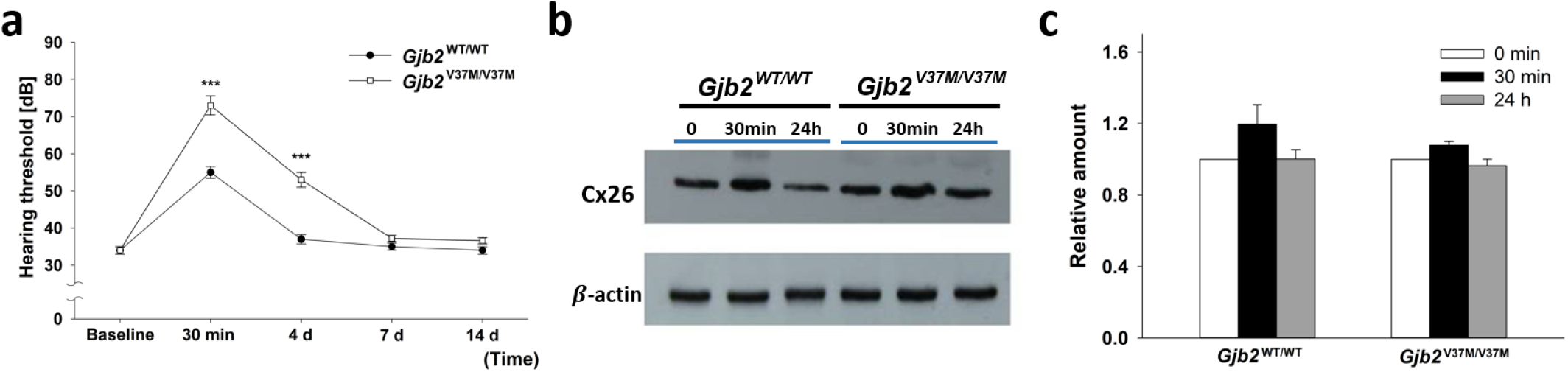
Higher vulnerability to noise in *Gjb2*^*V37M/V37M*^ mice. (a) *Gjb2*^*V37M/V37M*^ mice after 3 h of noise exposure revealed a significantly higher shift in hearing thresholds than *Gjb2*^*WT/WT*^ mice at 30 min and 4 d. (*n* = 5 for each strain; baseline: no noise exposure); (b) The western blot and (c) its relative amount of Cx26 expression after noise exposure (mean ± SEM) normalized to the counterpart before noise (0 min) in *Gjb2*^*WT/WT*^ and *Gjb2*^*V37M/V37M*^ mice, respectively. Both slightly increased immediately (30 min) but recovered to baseline level after 24 h without significant differences between the two groups.

To investigate the mechanisms underlying vulnerability to external noise, the expression of mCx26 was measured after noise exposure in the cochlea. The expression of mCx26 in the cochlea increased immediately after 30 min in both *Gjb2*^*WT/WT*^ and *Gjb2*^*V37M/V37M*^ mice but recovered to baseline levels after 24 h (Figure 4b). When normalized to the baseline Cx26 expression level before noise, there was no significant difference in the fold change of Cx26 expression at 30 min between *Gjb2*^*WT/WT*^ and *Gjb2*^*V37M/V37M*^ mice (1.19 ± 0.11 vs. 1.08 ± 0.02, Student’s t-test, *p* = 0.41; Figure 4c). These data indicate that the vulnerability of *Gjb2*^*V37M/V37M*^ mice to noise may be related to protein malfunction rather than decreased protein expression.

### 2.7 V37M resulted in aberrant interaction patterns between amino acid residues and blocked the mCx26 hemichannel

The blockage mechanisms caused by the missense V37M variant was elucidated using *in silico* model. Based on the reported crystal structures (PDBID: 2ZW3),^[23]^ Maeda *et al*. proposed that the gating mechanism of the Cx26 hemichannel was driven by a plug-mediated model with the plug (that is, the NTH domains) mainly stabilized by the Trp3-Met34 (W3-M34) sulfur-aromatic interaction. This interaction was ∼6 kJ/mol stronger than regular hydrophobic interactions^[28]^ and could make the NTHs adhere to the inner edge of the pore^[16c, 23]^ to maintain the open state of the pore. Several previous studies have argued that M34A and M34T variants would destabilize the affinity between NTHs and TM1, which would significantly disturb the open conformation of the hemichannel.^[16b, 16c]^

V37M, situated at the inner edge of the pore near M34, may spatially disturb the normal W3-M34 interaction. In our simulations with centralized vertical coordinates, the hexameric M37 (6-M37, hereinafter “hexameric” was abbreviated as “6-”) in V37M-mCx26 demonstrated downward shift and shorter distance from 6-W3 compared to the hexameric V37 (6-V37) in WT-mCx26 (**Figure 5**a-I and II). Both the 6-W3 and 6-M37 groups in V37M-mCx26 suggested a higher degree of aggregation than the WT (Figure 5a-III and IV). This result was rationalized by the affinity differences between the two pairs of hexameric groups, [6-Res37, 6-W3] and [6-M34, 6-W3] (stronger affinity corresponds to a larger negative quantity of binding free energy). Affinity of [6-V37, 6-W3] in WT-mCx26 was weaker than that of [6-M37, 6-W3] in V37M-mCx26 (−83.43 ± 0.16 vs. −98.25 ± 0.2 [kJ mol^-1^] in Figure 5a-V), whereas affinity of [6-M34, 6-W3] in WT-mCx26 was stronger than that of [6-M34, 6-W3] in V37M-mCx26 (−103.46 ± 0.17 vs. −94.72 ± 0.15 [kJ mol^-1^] in Figure 5a-IV). It could be inferred that 6-M37 competed with 6-M34 for the binding for 6-W3, resulting in a downward shift of 6-M37 that were “dragged” by 6-W3, tilting the inner edge to shrink the upper opening of the channel (Figure 5b), which was consistent with the reduced number of water molecules in V37M-Cx26 (Figure 1g). Collectively, our simulations indicate that the abnormal ion permeability of V37M-mCx26 could be attributed mainly to the increased affinity of M37 for the W3-containing NTHs, which caused the upper wall to tilt and collapse, resulting in pore shrinkage (Figure 4b).

**Figure 5.**
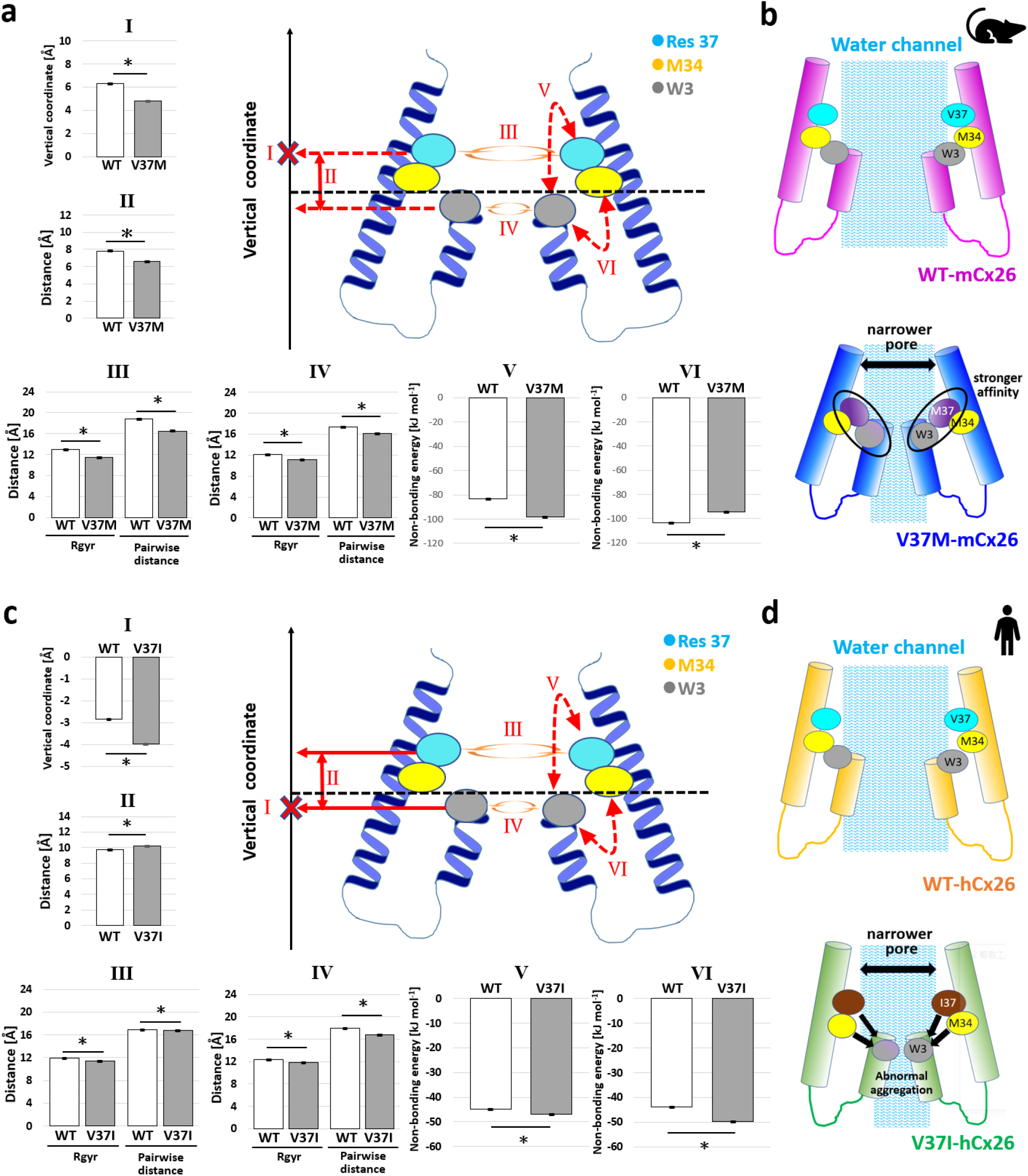
Characteristics of conformational changes and energetics derived from MD simulations to summarize the blockage mechanisms of the Cx26 hemichannel. (a and b) The computational quantities from the wild type and its mutant with homomeric V37M variants in mCx26. (c and d) The corresponding quantities from the wild type and its mutant with homomeric V37I variants in hCx26 protein. The vertical axis in panels a and c aligns with the membrane normal and meets the dash-lined horizontal axis at the membrane center (where 0 Å is at), defined as the averaged vertical coordinates of choline groups in both upper and lower leaflets of DOPC membrane; The “X” symbol marks the vertical coordinate of the target residue described in barplot I, which is the residue 37 and W3 in panel (a) and (c), respectively. All means ± SEM of computational quantities calculated from all snapshots of the equilibrated MD simulation systems are illustrated as bar plots with error bar. (*: *p* < 0.0001)

### 2.8 V37I caused abnormal NTHs aggregation and blocked the hCx26 hemichannel

The different interrupted ranges of water channels (Figure 1f and 1g) suggest different blockage mechanisms between V37I-hCx26 and V37M-mCx26. Our simulations revealed a downward shift of 6-W3 with increased distance from 6-Res37 in V37I-hCx26 (Figure 5c-I and -II), and both 6-I37 and 6-W3 in V37I-hCx26 demonstrated higher aggregation than those in WT-hCx26 (Figure 5c-III and -IV). The 6-W3 of V37I-hCx26 also revealed a similar affinity with 6-Res37 but a stronger affinity with 6-M34 compared to the counterparts of WT-hCx26 (Figure 5c-V and -VI). This indicated that 6-W3 would cause not only the blockage of the pore but also the proximity of the TM1 helices.

Accordingly, the abnormal shrinkage in V37I-hCx26 was initially attributed to the weaker affinity between I37 and NTHs, followed by the abnormal aggregation of W3-containing NTHs at the lower location of the hemichannel, and 6-W3 would further attract both 6-I37 and 6-M34 to form a narrower pore (Figure 5d), which interrupted the water channel (Figure 1f). The aggregation of NTHs also generated an environment unfavorable for the water molecules to stay in the pore and attracted the TM1 domain closer. Based collectively on these results, our *in silico* models indicate that the abnormal permeability of V37I-hCx26 was mainly attributed to abnormal NTHs aggregation caused by the reduced affinity with I37, which narrowed the pore volume and finally blocked the hemichannel.

## 3. Discussion

In this study, we established a MD simulation platform to investigate the structural dynamics of dysfunctional Cx26 hemichannels associated with *GJB2* variants. Our results deciphered the conformational changes of the Cx26 hemichannel conferred by the highly prevalent *GJB2* V37I variant in humans and successfully predicted the functional consequences and inheritance mode of the artificial *Gjb2* V37M variant in mice. Our MD simulations consistently demonstrated a narrow pore volume, reduced potassium permeability, and prolonged potassium ion retention time in both the V37I-hCx26 and V37M-mCx26 hemichannels. However, the blockage mechanisms of these two variants appeared different: the V37I variant in hCx26 caused abnormal aggregation of NTHs, leading to the subsequent shrunken pore volume and interrupted water channel, whereas the V37M variants in mCx26 resulted in aberrant binding patterns around M37 and tilted the inner edge of the hemichannel, followed by pore blockage (Figure 5).

Our MD simulations of V37I-hCx26 demonstrated a reduced number of water molecules and interruption at the lower part of the water channel (vertical coordinate ranging from 0 to −15 Å along the membrane normal shown in Figure 1f), which was highly correlated with the closer contact of NTHs (represented by the higher aggregation of 6-W3 in Figure 5c-IV). Therefore, it could be inferred that V37I would cause abnormal aggregation of NTHs promoted by a weakened affinity initially between 6-I37 and NTHs. This is followed by the attraction of TM1 domains closer to NTHs, which finally narrows the pore volume (Figure 5d). Furthermore, according to a previous study, the residue pair [Ile, Ile] had stronger inter-residue contact energies than the pair [Val, Val] (−5.69 vs. −4.81 in *RT* units for [Ile, Ile] and [Val, Val], respectively; *R*: gas constant (8.314 J K^−1^ mol^−1^); *T*: temperature (K)),^[29]^ which was in line with the higher aggregation of 6-Res37 in V37I-hCx26 than in WT-hCx26. By elucidating the conformational changes underlying the reduced permeability and conductance reported in previous *in vitro* biochemical studies,^[21]^ our MD simulations provide novel structural dynamics insights into the dysfunction of hCx26 caused by the V37I variant.

To verify the predictability of our MD simulation model in assessing the pathogenicity of other *GJB2* variants, an *in silico* mCx26 model was constructed with the artificial V37M variant and functional assays were performed using transgenic mice. Our MD simulations of homomeric V37M-mCx26 hemichannel demonstrated a reduced number of water molecules and an interrupted water channel around the M37 residue (vertical coordinates ranging from 0 to 10 Å along the membrane normal, shown in Figure 1g, which was highly correlated with the aggregation between 6-W3 and 6-M37. The conformational change was mainly attributed to the stronger affinity between 6-W3 and 6-Res37 in V37M-mCx26 than that in the wild-type mCx26, which was in line with the reported stronger inter-residue contact energies (−5.37 vs. −5.04 in *RT* units for [Met, Trp] and [Val, Trp], respectively).^[29]^ Consistent with these results, the subsequent animal studies showed that mice with the homozygous V37M variant (*Gjb2*^*V37M/V37M*^) developed hearing loss. Further functional experiments revealed that the expression and localization of Cx26 and GJs remained unchanged in homozygous mice; however, GJ-mediated metabolite transfer was affected by the V37M variant, indicating that the permeability of GJs was compromised in the cochlear sensory epithelium of homozygous mice.

Furthermore, our MD simulations demonstrated that heteromeric V37M-mCx26 did not compromise the functionality of the hemichannel, as the two predominant heteromeric V37M-mCx26 forms (that is, heteromer-1 and −3 in Figure 1d) retained normal pore volume and retention time of potassium compared to the WT-mCx26 hemichannel. Interestingly, this finding provides a plausible explanation for the normal hearing observed in mice with the heterozygous V37M variant (*Gjb2*^*WT/V37M*^): given that WT-mCx26 and V37M-mCx26 are equally expressed, the physiological function of GJs could be preserved. Notably, among the five mCx26 systems shown in Table 1, the 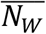 revealed high correlation with 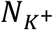 and 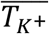, suggesting that compromised permeability was mainly attributed to the shrunken pore volume, and the three metrics could serve as reliable indicators to assess the dysfunction of Cx26 conferred by novel missense *GJB2*.

Interestingly, our animal studies revealed that mice homozygous for V37M are more vulnerable to noise exposure. These findings were consistent with those of a previous study that reported increased susceptibility to noise-induced hearing loss (NIHL) in *Gjb2* knockout mice.^[30]^ Several clinical studies have documented an association between NIHL and certain *GJB2* variants.^[31]^ It is likely that the increased susceptibility to NIHL in patients with specific *GJB2* genotypes, as demonstrated in the *Gjb2* knockout mice and our V37M knock-in mice, could be attributed to the compromised permeability of GJs to small metabolites or ions, such as potassium. Previous studies have revealed that potassium recycling in the cochlea may play a crucial role in preventing apoptosis of hair cells caused by NIHL.^[32]^ As such, these findings may have important clinical implications, in that noise exposure should be avoided in patients with specific *GJB2* variants.

The strength of this study lies in the comprehensive *in silico* demonstration of the blocking mechanism conferred by the human *GJB2* V37I variant through visualizing conformational changes in microsecond-scale simulations. Importantly, our subsequent *in intro* and *in vivo* experiments in mice with *Gjb2* V37M verified the predictive potential of our MD simulations for assessing the pathogenicity and inheritance modes of *GJB2* variants. However, some limitations of our MD simulation model merit further discussion. First, the MD simulation model focused only on the N-terminal and TM1 domains of Cx26. The sensitivity and reliability of conformational changes and pathogenicity of amino acid changes located in other domains of Cx26 require further validation. Second, our MD simulation model may not be applicable to assess missense variants with pathogenic mechanisms other than reduced permeability. This includes M1V, which results in aberrant translation;^[33]^ A88V, which results in mis-splicing at the RNA level;^[34]^ R32H and S199F, which result in mis-trafficking to form effective GJ plaques at the cell surface;^[35]^ and R184P, which results in reduced oligomerization.^[33]^

It has been reported that *GJB2* variants can lead to SNHI through digenic inheritance with *GJB6*^[36]^ and *GJB3* variants^[37]^, respectively; in parallel with the clinical findings, Cx26 can oligomerize with Cx30 and Cx31 to form heteromeric or heterotypic Cx26/Cx30^[38]^ and Cx26/Cx31^[39]^ channels in the cochlea. In principle, each Cx26 monomer in our model could be freely replaced with isoforms of other SNHI-related connexins to co-assemble a heteromeric hemichannel or a heterotypic GJ channel. Unfortunately, the crystal or cryo-electron microscopy structures for most SNHI-related connexins remain unsolved for the time being. Putative structures constructed using homology modeling^[40]^ or AlphaFold^[41]^ may enable the generation of heteromeric/heterotypic channels for investigating digenic pathogenetics in the future.

In conclusion, we revealed the channel closure mechanism of the highly prevalent *GJB2* V37I variant, and the functional consequences as well as the inheritance pattern of the tested *Gjb2* V37M variant in mice were cogently predicted. Our MD simulations demonstrated that both variants led to a narrowed pore volume, reduced potassium permeability, and prolonged potassium ion retention time through different blockage mechanisms. As a proof-of-concept, our MD simulations could be developed as an assessment tool for addressing the pathogenesis and inheritance of *GJB2*-related SNHI as well as other diseases caused by connexin dysfunction.

## 4. Experimental Section

### Construction of the membrane-embedded Cx26 model in silico

The original tertiary crystal structure of human connexin 26 (Cx26) protein^[23]^ (PDB ID: 2ZW3) had missing residues (residues 110–124) and N-terminal amino acid residue (Met1). We used SWISS-MODEL^[24]^ and open-source software PyMOL (Version 2.3.3, Schrödinger, LLC)^[42]^ to structurally model the missing loop and Met1 based on the primary sequence of human Cx26 protein (UniProt ID: P29033), while the hexameric hemichannel was constructed by superimposing the single-chain homology model onto each chain of the crystal structure (PDB ID: 2ZW3). The protonation state of protein in this wild-type human Cx26 hemichannel (WT-hCx26) at neutral pH was suggested by PROPKA^[43]^ (Version 3.1). Next, WT-hCx26 embedded in a phospholipid bilayer composed of 444 1,2-dioleoyl-*sn*-glycero-3-phosphocholine (DOPC) molecules in a 14 × 14 × 19 [nm^3^] rectangular box was generated using CHARMM-GUI,^[44]^ which was solvated in bulk non-polar water molecules with 150 mM KCl to mimic a high potassium concentration in the endolymph of the scala media.^[45]^ A protein mutant with the homomeric variant V37I (V37I-hCx26) was constructed based on this WT-hCx26 model using the same pipeline.

In a similar vein, using SWISS-MODEL,^[24]^ a structural model of mouse connexin 26 (mCx26) and its V37M variant were constructed using mouse Cx26 protein sequence (UniProt ID: Q00977) and the crystal structure of human connexin 26 (Cx26) protein (PDB ID: 2ZW3).^[23]^

### Molecular dynamics (MD) simulations

All simulations (∼30 μs in total) were performed using the GROMACS package program^[46]^ and Martini force field.^[47]^ The pressure and temperature were maintained at 1 bar and 310 K, respectively, and the cut-off values for both Coulomb and Van der Waals forces was 1.1 nm. All simulations were performed for 4 μs in NPT ensemble (i.e., simulation in isothermal-isobaric ensemble maintaining constant particle *N*, pressure *P*, and temperature *T*) controlled by Parrinello–Rahman barostat and Berendsen thermostat after at least 30 ns of equilibration. All trajectories of the MD simulations were calculated at 20 fs per step, and the coordinates were recorded every 1 ns. Subsequently, the vertical coordinates of the rectangular periodic boundary box per frame were centered on the mean vertical coordinates of all the choline groups in both the upper and lower leaflets.

### Determination of pore volume, potassium permeability, and retention time of potassium ions

The pore volume of Cx26 hemichannel was represented by the average number of water molecules 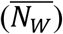 inside the hemichannel pore per snapshot (10 ns per frame in a 4 μs simulation) and within the average vertical coordinates of DOPC beads located in both the upper and lower leaflets. The potassium permeability 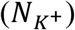 was calculated as the total number of potassium ions passing through the pore of the hemichannel that was determined according to the trajectory at which they entered from either entrance and finally left from the opposite gate (see Supplementary Video S1 and S3 as examples) during the 4 μs simulation. The retention time of potassium 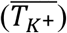 was obtained by averaging the time of all potassium ions that stay inside the channel for at least 20 ns where ions are considered inside the channel when their vertical coordinates are in between the average positions of choline beads in the upper and lower DOPC leaflets.

### Generation of knock-in mice

A mutant gene-targeting vector was constructed using recombineering approach, previously developed by Copeland et al.^[48]^ A BAC clone (clone no. bMQ369p05 Geneservice™) from 129S7/AB2.2 BAC library, containing the mouse *Gjb2* genomic region, was used to construct the gene-targeting vector (**Figure S1**a). The BAC was transferred to the modified *E. coli* strain, EL350, by electroporation. The subcloned 15.9 kb genomic region was modified in a subsequent targeting round by inserting a neomycin (*neo*) cassette from PL452 plasmid and creating the V37M variant. The conditional targeting vector was linearized by *Not*I digestion and electroporated into R1 embryonic stem (ES) cells. G418 (240 μg mL^−1^) and ganciclovir (2 μM) double-resistant clones were analyzed using Southern blot hybridization (Figure S1b). The retained *neo* cassette flanked by *loxP* sites was excised *in vivo* by transfection of the targeted clone with a plasmid transiently expressing Cre recombinase. The established ES clones were identified by PCR screening and subsequently injected into C57BL/6 blastocysts to produce chimeras. The mice were maintained on a C57BL/6 genetic background for further experiments. Heterozygous mice were bred to obtain *Gjb2*^*WT/WT*^, *Gjb2*^*V37M/WT*^, and *Gjb2*^*V37M/V37M*^ mice (Figure S1c). All animal experiments were carried out in accordance with the animal welfare guidelines approved by the National Taiwan University Hospital (approval no. 200700204).

### Audiological measurements

Mice were anesthetized with sodium pentobarbital (35 mg kg^−1^), delivered intraperitoneally and maintained in a head-holder within an acoustically and electrically insulated and grounded test room. An evoked potential detection system (Smart EP 3.90, Intelligent Hearing Systems, Miami, FL) was used to measure the auditory brainstem response (ABR) thresholds in mice. Click sounds as well as 8, 16, and 32 kHz tone bursts at varying intensities (from 10 dB to 130 dB SPL), were given to evoke ABRs in mice.^[27]^

### Inner ear morphology

The inner ear tissues of mice were subjected to hematoxylin and eosin (H&E) staining and whole-mount studies.^[49]^ The H&E-stained tissues were examined using a Nikon Optiphot-2 microscope. For whole-mount studies, the tissues were stained with rhodamine–phalloidin (1:100 dilution; Molecular Probes, Eugene, OR, USA) and images were obtained using a laser scanning confocal microscope (Zeiss LSM 510, Germany).

### Protein expression

Flattened cochlear preparations^[26]^ obtained from P8 mice were fixed in paraformaldehyde (4%), permeabilized in Triton X-100 (0.5%), and blocked in goat serum (10%). After incubation with primary antibodies against Cx26 (Zymed Labs, South San Francisco, CA, USA) and FITC-conjugated goat anti-rabbit secondary antibodies, the samples were examined using a laser scanning confocal microscope (Zeiss LSM 510, Carl Zeiss, Germany). The fluorescence intensity of Cx26 immunoreactivities was quantified using Metamorph 7.1 software (Universal Imaging Corporation, Downingtown, PA) in three *Gjb2*^*WT/WT*^ and three *Gjb2*^*V37M/V37M*^ mice. For the inner ear sample of each animal, five images were randomly selected, and the fluorescence intensity of the images was calculated and compared between that of the *Gjb2*^*WT/WT*^ and *Gjb2*^*V37M/V37M*^ mice.

### Fluorescent dye diffusion

Flattened cochlear preparations^[26]^ obtained from P8 mice (C57BL/6 background) were placed in a recording chamber mounted on an upright microscope (Axioskop2 FX plus; Carl Zeiss, Germany). An extracellular solution (HBSS) was perfused at a rate of 1 drop s^−1^. DIC optics enabled us to directly recognize different cochlear cell types in the acutely isolated cochlear segment. Membrane-impermeable fluorescent dyes were injected into the cytoplasm of a single cell by forming the whole-cell patch-clamp recording mode that was confirmed by monitoring the whole-cell resistance and capacitance using a standard protocol (Axonpatch 200 B, Axon Instruments, CA). For the preparation of fluorescent dyes, PI (0.75 mM, MW= 668 Da, charge = +2, catalog no. P1304MP), and LY (1 mM, MW=457 Da, charge = −2, catalog no. L-453) were added to the pipette solution containing KCl (120 mM), MgCl_2_ (1 mM), HEPES (10 mM), and EGTA (10 mM). Intercellular dye diffusion patterns were recorded using a digital single-lens reflex (D700, Nikon, Japan) at various time points after the establishment of the whole-cell recording conformer.

### Noise exposure experiments

Twelve-week-old mice were exposed to octave band noise for 3 h with a Bruel & Kjaer 2238 sound level meter peak at 4 kHz and 115 dB SPL.^[25]^ The noise room was fitted with a speaker (model NO. 1700-2002, Grason-Stadler Inc., MN) driven by a noise generator (GSI-61, Grason-Stadler Inc., MN) and power amplifier (5507-Power, TECHRON, Union City, IN). After 3h noise exposure, ABR thresholds at clicks were then recorded in mice after 30 min and 4, 7, and 14 d.

In addition to hearing levels, Cx26 expression after the noise exposure was also investigated. Protein extracts of the inner ear were supplemented with DTT, heated for 5 min at 95 °C, separated by SDS gel electrophoresis, and transferred to a PVDF membrane (Amersham Pharmacia Biotech, Little Chalfont, UK) by semi-dry electroblotting. The PVDF membranes were developed using an enhanced chemiluminescence western blot detection kit (Pierce SuperSignal® West Dura, Rockford, IL) and exposed to Lumi-Film chemiluminescent detection films (Roche Diagnostics, Mannheim, Germany). The experiment was replicated five times, and the results were averaged.

### Analyses of affinities between pairs of hexameric residue/NTHs clusters

The affinity calculated between two targeted hexameric residue clusters was determined by non-bonding energies composed of both electrostatic and van der Waals forces based on the Martini force fields. Each hexameric residue cluster was defined as a group of six residues from each monomer of the hexameric Cx26 hemichannel. The affinities were recorded every 1 ns during the 4 μs simulation to calculate the mean ± SEM.

### Assessment of the degree of aggregation in the Cx26 hemichannel

Two physical quantities were calculated to assess the degree of aggregation: average pairwise distance and radius of gyration (*R*_*gyr*_). The pairwise distance was determined by recording the average residue-residue distance within each hexameric residue cluster (i.e., the average distances of a total of 15 pairs of residues for each hexameric residue cluster) per snapshot (1 ns per frame) during the 4 μs simulation and represented as mean ± SEM. The *R*_*gyr*_ was determined by recording all 3D coordinates of residues within each hexameric residue cluster to calculate *R*_*gyr*_ per snapshot during the 4 μs simulation and represented as mean ± SEM. The *R*_*gyr*_ equation is as follows:

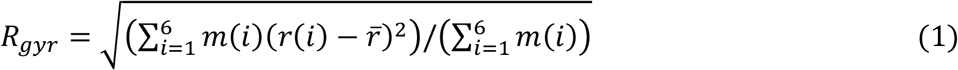

where *r(i)* and *m(i)* are the position and molecular mass of the *i*^th^ residue within each hexameric residue cluster, respectively, and 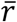 is the weighted average position per hexameric residue cluster.

## Supporting Information

Supporting Information is available from the Wiley Online Library or from the author.

## Acknowledgments

We would like to thank technical support from the Taiwan Mouse Clinic and Transgenic Mouse Models Core Facility of National Core Facility Program for Biotechnology supported by the Ministry of Science and Technology.

## Conflict of interest

The authors declare no conflict of interest

## Data availability statement

The data that support the findings of this study are available from the corresponding author upon reasonable request.

## Funding statement

This study was supported by research grants from National Health Research Institutes (NHRI-EX111-10914PI), as well as the collaborative project between National Tsing Hua University and National Taiwan University Hospital Hsin-Chu Branch (110Q2528E1).

## Notes

### Competing Interest Statement

The authors have declared no competing interest.

## References

[1] J. C. Wingard, H.-B. Zhao, Frontiers in Cellular Neuroscience 2015, 9, 202.

[2] a)A. Kenneson, K. Van Naarden Braun, C. Boyle, Genetics in Medicine 2002, 4 (4), 258; b)R. L. Snoeckx, P. L. Huygen, D. Feldmann, S. Marlin, F. Denoyelle, J. Waligora, M. Mueller-Malesinska, A. Pollak, R. Ploski, A. Murgia, The American Journal of Human Genetics 2005, 77 (6), 945.

[3] a)T. Kikuchi, R. S. Kimura, D. L. Paul, J. C. Adams, Anatomy and embryology 1995, 191 (2), 101; b)A. Forge, D. Becker, S. Casalotti, J. Edwards, N. Marziano, G. Nevill, Journal of Comparative Neurology 2003, 467 (2), 207; c)H. B. Zhao, N. Yu, Journal of Comparative Neurology 2006, 499 (3), 506.

[4] a)D. P. Kelsell, J. Dunlop, H. P. Stevens, N. J. Lench, J. Liang, G. Parry, R. F. Mueller, I. M. Leigh, Nature 1997, 387 (6628), 80; b)P. Wangemann, Hearing research 2002, 165 (1-2), 1.

[5] M. Beltramello, V. Piazza, F. F. Bukauskas, T. Pozzan, F. Mammano, Nature cell biology 2005, 7 (1), 63.

[6] a)Q. Chang, W. Tang, Y. Kim, X. Lin, Neurobiology of disease 2015, 73, 418; b)Y. Wang, Q. Chang, W. Tang, Y. Sun, B. Zhou, H. Li, X. Lin, Biochemical and biophysical research communications 2009, 385 (1), 33.

[7] H. Azaiez, K. T. Booth, S. S. Ephraim, B. Crone, E. A. Black-Ziegelbein, R. J. Marini, A. E. Shearer, C. M. Sloan-Heggen, D. Kolbe, T. Casavant, The American Journal of Human Genetics 2018, 103 (4), 484.

[8] Q. Wang, E. Pierce-Hoffman, B. B. Cummings, J. Alföldi, L. C. Francioli, L. D. Gauthier, A. J. Hill, A.H. O’Donnell-Luria, K. J. Karczewski, D. G. MacArthur, Nature communications 2020, 11 (1), 1.

[9] S. Abe, S.-i. Usami, H. Shinkawa, P. M. Kelley, W. J. Kimberling, Journal of medical genetics 2000, 37 (1), 41.

[10] S. Y. Kim, G. Park, K.-H. Han, A. Kim, J.-W. Koo, S. O. Chang, S. H. Oh, W.-Y. Park, B. Y. Choi, PloS one 2013, 8 (4), e61592.

[11] D. Wattanasirichaigoon, C. Limwongse, C. Jariengprasert, P. Yenchitsomanus, C. Tocharoenthanaphol, W. Thongnoppakhun, C. Thawil, D. Charoenpipop, T. Pho-Iam, S. Thongpradit, Clinical genetics 2004, 66 (5), 452.

[12] a)L. Li, J. Lu, Z. Tao, Q. Huang, Y. Chai, X. Li, Z. Huang, Y. Li, M. Xiang, J. Yang, PloS one 2012, 7 (5), e36621; b)H.-L. Hwa, T.-M. Ko, C.-J. Hsu, C.-H. Huang, Y.-L. Chiang, J.-L. Oong, C.-C. Chen, C.-K. Hsu, Genetics in Medicine 2003, 5 (3), 161.

[13] Y. Chai, D. Chen, L. Sun, L. Li, Y. Chen, X. Pang, L. Zhang, H. Wu, T. Yang, Clinical genetics 2015, 87 (4), 350.

[14] a)A. Pollak, A. Skórka, M. Mueller-Malesińska, G. Kostrzewa, B. Kisiel, J. Waligóra, P. Krajewski, M. Ołdak, L. Korniszewski, H. Skarżyński, American journal of medical genetics Part A 2007, 143 (21), 2534; b)K. Tsukada, S. Nishio, S. Usami, D. G. S. Consortium, Clinical genetics 2010, 78 (5), 464; c)S. Huang, B. Huang, G. Wang, Y. Yuan, P. Dai, PLoS One 2015, 10 (6), e0129662; d)E. Gallant, L. Francey, E. A. Tsai, M. Berman, Y. Zhao, H. Fetting, M. Kaur, M. A. Deardorff, A. Wilkens, D. Clark, American Journal of Medical Genetics Part A 2013, 161 (9), 2148; e)M. A. Kenna, H. A. Feldman, M. W. Neault, A. Frangulov, B.-L. Wu, B. Fligor, H. L. Rehm, Archives of Otolaryngology–Head & Neck Surgery 2010, 136 (1), 81.

[15] a)C.-C. Wu, C.-H. Tsai, C.-C. Hung, Y.-H. Lin, Y.-H. Lin, F.-L. Huang, P.-N. Tsao, Y.-N. Su, Y. L. Lee, W.-S. Hsieh, Genetics in Medicine 2017, 19 (1), 6; b)P.-Y. Chen, Y.-H. Lin, T.-C. Liu, Y.-H. Lin, L.-H. Tseng, T.-H. Yang, P.-L. Chen, C.-C. Wu, C.-J. Hsu, Ear and Hearing 2020, 41 (1), 143.

[16] a)O. Jara, R. Acuña, I. E. García, J. Maripillán, V. Figueroa, J. C. Sáez, R. Araya-Secchi, C. F. Lagos, T. Pérez-Acle, V. M. Berthoud, Molecular biology of the cell 2012, 23 (17), 3299; b)A. Oshima, K. Tani, M. M. Toloue, Y. Hiroaki, A. Smock, S. Inukai, A. Cone, B. J. Nicholson, G. E. Sosinsky, Y. Fujiyoshi, Journal of molecular biology 2011, 405 (3), 724; c)F. Zonta, D. Buratto, C. Cassini, M. Bortolozzi, F. Mammano, Frontiers in physiology 2014, 5, 85.

[17] a)P. E. Purnick, D. C. Benjamin, V. K. Verselis, T. A. Bargiello, T. L. Dowd, Archives of biochemistry and biophysics 2000, 381 (2), 181; b)A. Oshima, T. Doi, K. Mitsuoka, S. Maeda, Y. Fujiyoshi, Journal of Biological Chemistry 2003, 278 (3), 1807; c)A. Oshima, FEBS letters 2014, 588 (8), 1230.

[18] A. Batool, S. Yasmeen, S. Rashid, Journal of Molecular Liquids 2017, 227, 168.

[19] I. E. García, F. Villanelo, G. F. Contreras, A. Pupo, B. I. Pinto, J. E. Contreras, T. Pérez-Acle, O. Alvarez, R. Latorre, A.D. Martínez, Journal of General Physiology 2018, 150 (5), 697.

[20] J. M. Valdez Capuccino, P. Chatterjee, I. E. García, W. M. Botello-Smith, H. Zhang, A. L. Harris, Y. Luo, J. E. Contreras, Journal of General Physiology 2019, 151 (3), 328.

[21] a)R. Bruzzone, V. Veronesi, D. Gomes, M. Bicego, N. Duval, S. Marlin, C. Petit, P. D’Andrea, T. White, FEBS letters 2003, 533, 79; b)M. Palmada, K. Schmalisch, C. Böhmer, N. Schug, M. Pfister, F. Lang, N. Blin, Neurobiology of disease 2006, 22 (1), 112; c)J. Kim, J. Jung, M. G. Lee, J. Y. Choi, K.-A. Lee, Experimental & molecular medicine 2015, 47 (6), e169.

[22] F. J. Del Castillo, I. Del Castillo, Frontiers in molecular neuroscience 2017, 10, 428.

[23] S. Maeda, S. Nakagawa, M. Suga, E. Yamashita, A. Oshima, Y. Fujiyoshi, T. Tsukihara, Nature 2009, 458 (7238), 597.

[24] M. Biasini, S. Bienert, A. Waterhouse, K. Arnold, G. Studer, T. Schmidt, F. Kiefer, T. G. Cassarino, M. Bertoni, L. Bordoli, Nucleic acids research 2014, 42 (W1), W252.

[25] Q. Y. Zheng, K. R. Johnson, L. C. Erway, Hearing research 1999, 130 (1-2), 94.

[26] Q. Chang, W. Tang, S. Ahmad, B. Zhou, X. Lin, PLoS One 2008, 3 (12), e4088.

[27] Y.-C. Lu, C.-C. Wu, T.-H. Yang, Y.-H. Lin, I.-S. Yu, S.-W. Lin, Q. Chang, X. Lin, J.-M. Wong, C.-J. Hsu, PloS one 2013, 8 (6), e64906.

[28] C. C. Valley, A. Cembran, J. D. Perlmutter, A. K. Lewis, N. P. Labello, J. Gao, J. N. Sachs, Journal of Biological Chemistry 2012, 287 (42), 34979.

[29] I. Bahar, R. L. Jernigan, Journal of molecular biology 1997, 266 (1), 195.

[30] X.-X. Zhou, S. Chen, L. Xie, Y.-Z. Ji, X. Wu, W.-W. Wang, Q. Yang, J.-T. Yu, Y. Sun, X. Lin, International journal of molecular sciences 2016, 17 (3), 301.

[31] a)M. Pawelczyk, L. Van Laer, E. Fransen, E. Rajkowska, A. Konings, P. I. Carlsson, E. Borg, G. Van Camp, M. Sliwinska-Kowalska, Annals of human genetics 2009, 73 (4), 411; b)W. S. Li, Y. L. Gang, L. R. Ping, Z. W. Zhan, G. W. Min, X. L. Ping, J. Xu, Z. Y. Han, Y. Ding, C. Dong, Biomedical and environmental sciences 2014, 27 (12), 965.

[32] a)A. A. Zdebik, P. Wangemann, T. J. Jentsch, Physiology 2009, 24 (5), 307; b)M. Cohen-Salmon, T. Ott, V. Michel, J.-P. Hardelin, I. Perfettini, M. Eybalin, T. Wu, D. C. Marcus, P. Wangemann, K. Willecke, Current Biology 2002, 12 (13), 1106.

[33] E. Thönnissen, R. Rabionet, M. Arbonès, X. Estivill, K. Willecke, T. Ott, Human genetics 2002, 111 (2), 190.

[34] J. Cook, E. de Wolf, N. Dale, Royal Society open science 2019, 6 (8), 191128.

[35] Z. Xiao, Z. Yang, X. Liu, D. Xie, Acta oto-laryngologica 2011, 131 (1), 59.

[36] E. Cama, S. Melchionda, T. Palladino, M. Carella, R. Santarelli, E. Genovese, F. Benettazzo, L. Zelante, E. Arslan, International journal of audiology 2009, 48 (1), 12.

[37] K. Chen, X. Wu, L. Zong, H. Jiang, Journal of clinical laboratory analysis 2018, 32 (9), e22592.

[38] J. Defourny, N. Thelen, M. Thiry, Mechanisms of Development 2019, 155, 8.

[39] X.-Z. Liu, Y. Yuan, D. Yan, E. H. Ding, X. M. Ouyang, Y. Fei, W. Tang, H. Yuan, Q. Chang, L. L. Du, Human genetics 2009, 125 (1), 53.

[40] a)J. Yang, Y. Zhang, Current protocols in bioinformatics 2015, 52 (1), 5.8. 1; b)A. Waterhouse, M. Bertoni, S. Bienert, G. Studer, G. Tauriello, R. Gumienny, F. T. Heer, T. A. P. de Beer, C. Rempfer, L. Bordoli, Nucleic acids research 2018, 46 (W1), W296.

[41] J. Jumper, R. Evans, A. Pritzel, T. Green, M. Figurnov, O. Ronneberger, K. Tunyasuvunakool, R. Bates, A. Žídek, A. Potapenko, Nature 2021, 596 (7873), 583.

[42] M. A. Lill, M. L. Danielson, Journal of computer-aided molecular design 2011, 25 (1), 13.

[43] M. H. Olsson, C. R. Søndergaard, M. Rostkowski, J. H. Jensen, Journal of chemical theory and computation 2011, 7 (2), 525.

[44] a)S. Jo, T. Kim, V. G. Iyer, W. Im, Journal of computational chemistry 2008, 29 (11), 1859; b)Y. Qi, H. I. Ingólfsson, X. Cheng, J. Lee, S. J. Marrink, W. Im, Journal of chemical theory and computation 2015, 11 (9), 4486.

[45] H. Hibino, Y. Kurachi, Physiology 2006, 21 (5), 336.

[46] M. J. Abraham, T. Murtola, R. Schulz, S. Páll, J. C. Smith, B. Hess, E. Lindahl, SoftwareX 2015, 1, 19.

[47] a)D. H. de Jong, G. Singh, W. D. Bennett, C. Arnarez, T. A. Wassenaar, L. V. Schafer, X. Periole, D. P. Tieleman, S. J. Marrink, Journal of chemical theory and computation 2013, 9 (1), 687; b)T. A. Wassenaar, H. I. Ingólfsson, R. A. Bockmann, D. P. Tieleman, S. J. Marrink, Journal of chemical theory and computation 2015, 11 (5), 2144.

[48] a)E.-C. Lee, D. Yu, J. M. De Velasco, L. Tessarollo, D. A. Swing, D. L. Court, N. A. Jenkins, N. G. Copeland, Genomics 2001, 73 (1), 56; b)K.-Y. Su, W.-L. Chien, W.-M. Fu, I.-S. Yu, H.-P. Huang, P.-H. Huang, S.-R. Lin, J.-Y. Shih, Y.-L. Lin, Y.-P. Hsueh, Journal of Neuroscience 2007, 27 (10), 2513.

[49] Y.-C. Lu, C.-C. Wu, W.-S. Shen, T.-H. Yang, T.-H. Yeh, P.-J. Chen, I.-S. Yu, S.-W. Lin, J.-M. Wong, Q. Chang, PLoS One 2011, 6 (7), e22150.

