## Supplementary Material for "Computer simulations reveal pathogenicity and inheritance modes of hearing loss-causing germinal variants"

**Supplementary Video 1**

The potassium ions successfully pass through the pore of WT-hCx26.

**Supplementary Video 2**

The potassium ions fail to pass through the pore of V37I-hCx26.

**Supplementary Video 3**

The potassium ions successfully pass through the pore of WT-mCx26.

**Supplementary Video 4**

The potassium ions fail to pass through the pore of V37M-mCx26.

**Table S1. Hearing thresholds of knock-in mice for wild-type, heterozygous, and homozygous V37M at any frequency and age measured by their auditory brainstem response**

| Frequency | Time (wks) | ***Gjb2*** $^{WT/WT}$ | ***Gjb2*** $^{WT/V37M}$ | ***Gjb2*** $^{V37M/V37M}$ |
|---|---|---|---|---|
| click | 4 | 35.38 ± 1.05 [a)] | 35.26 ± 0.81 | 36.00 ± 1.06 |
| | 12 | 33.85 ± 1.40 | 37.11 ± 1.29 | 39.00 ± 1.23 * [b)] |
| | 28 | 37.69 ± 0.92 | 37.89 ± 0.88 | 46.17 ± 1.33 *** |
| | 44 | 37.31 ± 2.51 | 40.26 ± 1.81 | 56.33 ± 2.34 *** |
| 8k | 4 | 39.23 ± 1.11 | 39.47 ± 1.07 | 39.17 ± 0.99 |
| | 12 | 41.54 ± 1.19 | 42.11 ± 1.17 | 42.00 ± 1.24 |
| | 28 | 40.38 ± 1.74 | 41.05 ± 1.24 | 47.00 ± 1.26 ** |
| | 44 | 45.00 ± 2.12 | 45.79 ± 2.37 | 55.83 ± 1.56 *** |
| 16k | 4 | 22.69 ± 1.66 | 23.68 ± 0.64 | 23.33 ± 0.81 |
| | 12 | 27.31 ± 1.08 | 25.79 ± 1.22 | 28.83 ± 1.21 |
| | 28 | 27.31 ± 2.09 | 28.16 ± 1.03 | 38.00 ± 1.45 *** |
| | 44 | 36.15 ± 2.66 | 35.26 ± 2.24 | 47.50 ± 2.18 ** |
| 32k | 4 | 25.77 ± 2.46 | 27.11 ± 2.07 | 28.50 ± 0.80 |
| | 12 | 29.23 ± 2.39 | 29.47 ± 1.66 | 34.00 ± 1.08 * |
| | 28 | 29.23 ± 2.32 | 30.79 ± 1.49 | 40.17 ± 1.21 *** |
| | 44 | 35.38 ± 2.37 | 36.05 ± 1.90 | 54.17 ± 2.48 *** |

[a)] Hearing threshold (mean ± SEM) in dB using sound pressure level (SPL); [b)] p-value compared to the wild type (***, $p < 0.001$; **, $p < 0.01$; *, $p < 0.05$).

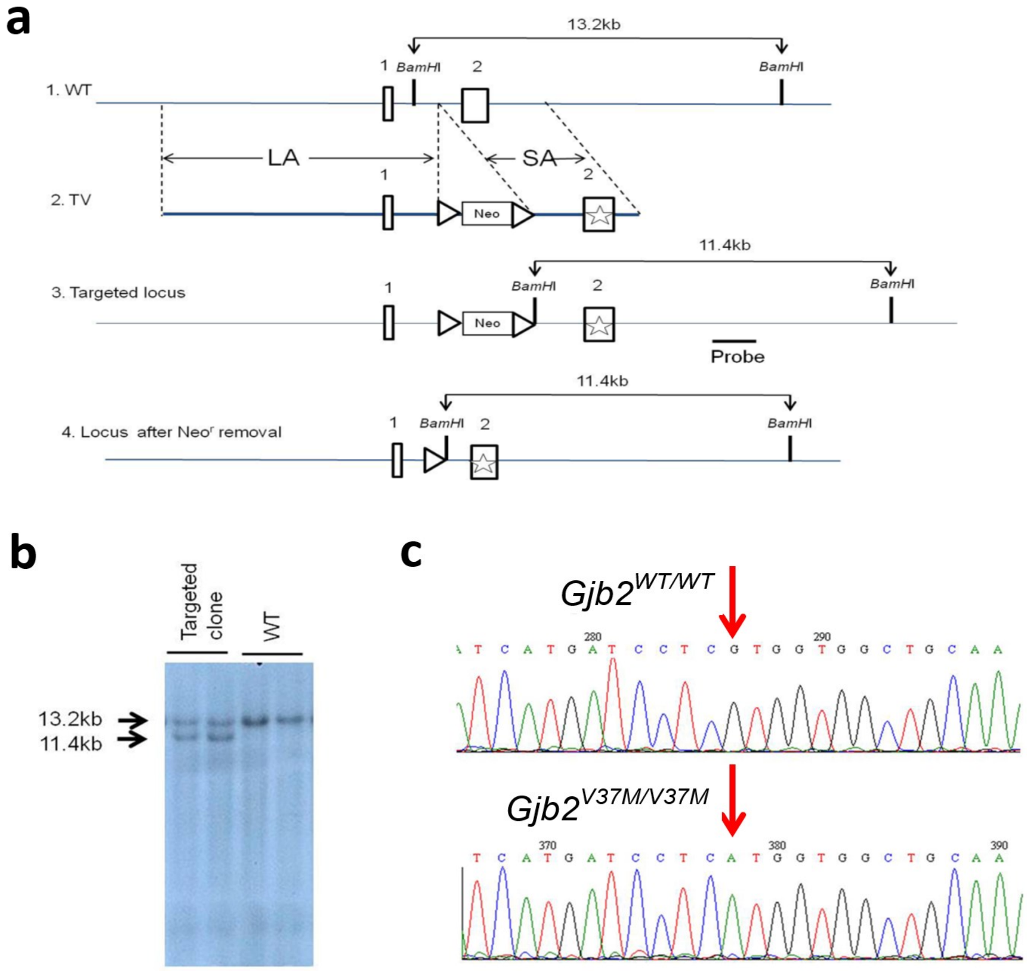


**Figure S1.** Generation of mice with the *Gjb2* p.V37M (c.109G>A) variant.
